# Conserved and diverged mechanisms for RNA target-site recognition by the multi KH-domain Bicaudal C protein

**DOI:** 10.1101/2022.07.22.501140

**Authors:** Megan E. Dowdle, Tommaso Pietro Tonelli, Catherine A. Fox, Michael D. Sheets

## Abstract

The Bicaudal-C (Bicc1) multi-KH domain RNA binding protein controls cell fates by binding and repressing the translation of specific target mRNAs. To provide new insights into Bicc1-RNA binding we used quantitative assays to analyze interactions between *Xenopus* Bicc1 protein and the 32-nt Bicc1 RNA target site from *Xl*Cripto-1 mRNA. High-affinity binding by *Xl*Bicc1(K_d_=34nM) required both a 5’ leader sequence and a 3’loop sequence. Bicc1 proteins from humans (*Hs*Bicc1), zebrafish (*Dr*Bicc1) and fruit flies (*Dm*Bicc1) also bound the *Xl*Cripto-1 RNA with high-affinity via an interaction requiring the 5’ leader sequence, providing evidence for a conserved mechanism of RNA recognition by Bicc1. However, in contrast to *Xl*Bicc1, each of the other species’ Bicc1s showed a significantly relaxed requirement for the 3’ loop sequence. Analyses of mutant forms of *Xl*Bicc1, revealed that the KH1 domain promoted Bicc1’s requirement for the loop sequence via a GXXG-independent mechanism. In addition, a fusion protein comprised of the *Xl*KH1 and the *Dm*Bicc1-KH2-KH3 subdomains was sufficient to establish high-affinity, loop-sensitive Bicc1-binding to *Xl*Cripto-1 RNA. These data support a model wherein the KH2 domain interacts with the leader sequence via a GXXG-mediated KH-RNA interface, defining the conserved core of the metazoan Bicc1-RNA interaction, while the KH1 domain is required for the formation of a more flexible, secondary protein-RNA interface that allows recognition of the 3’ loop sequence. This secondary interface differs between species and possibly between target mRNAs, establishing the potential for a range of translational repression activities for Bicc1 target mRNAs.

## INTRODUCTION

Bicaudal-C (Bicc1) is a conserved RNA binding protein that regulates metazoan development by repressing the translation of specific mRNAs (Mahone et al. 1995; Gamberi and Lasko 2012). Bicc1-mediated repression controls key developmental decisions. For example, during the maternal stages of *Drosophila* and *Xenopus* embryogenesis, defects in Bicc1 lead to aberrant anterior-posterior patterning (Bull 1966; Mohler and Wieschaus 1985, 1986; Wessely and De Robertis 2000; Park et al. 2016). At later stages of development, Bicc1 influences Left/Right patterning and proper organ formation and function (Minegishi et al. 2021; Maerker et al. 2021). Despite Bicc1’s biological relevance, the mechanisms by which its selects mRNA targets for repression are not clear.

Studies of *Xenopus* Bicc1 (*Xl*Bicc1) provide insights into Bicc1-dependent mechanisms of selective RNA binding and repression (Zhang et al. 2013, 2014; Dowdle et al. 2019). The N-terminal region of Bicc1 contains three canonical KH-domains and two KH-like (KHL) domains in the following order: KH1-KH2-KHL1-KH3-KHL2, which will be referred to as KH1-KH3. The paradigmatic KH domain-RNA interface involves a conserved GXXG motif, and recognition of 3-4 nucleotides (Hollingworth et al. 2012; Nicastro et al. 2015, 2017). Less is known about the RNA-protein interfaces involving multi-KH domain containing proteins, but their RNA target sites appear more complex. For example, the minimal high-affinity and functional *Xl*Bicc1 target site within the Cripto-1 mRNA’s 3’ UTR is 32-nt and contains a 5’ leader sequence followed by a putative 3’ stem-loop structure, suggestive of a considerably more complex RNA-protein interface than predicted for single KH-domain-RNA interactions (Zhang et al. 2014). In support of this idea, experimental analyses reveal that both RNA sequence and structural features of the Cripto-1 RNA site are important for Bicc1 recognition (Zhang et al. 2014). Furthermore, each of the KH domains within the Bicc1 RNA binding domain is essential for Cripto-1 RNA binding, but only the conserved GXXG motif of the most conserved KH domain, the KH2 domain, is required (Dowdle et al. 2019). Thus the additional KH domains must directly and/or indirectly provide for an additional non-paradigmatic (i.e. GXXG-independent) RNA binding surface.

In this study, the strengths of multiple distinct wild-type and variant Bicc1-Cripto-1 RNA interfaces were determined using fluorescence polarization to detect the formation of Bicc1-RNA complexes. The data reveal that the *Xl*Bicc1 binds to the defined *Xl*Cripto-RNA Bicc1 site via an interaction requiring at least two distinct protein-RNA interfaces. The first protein-RNA interface requires the GXXG motif within the KH2 domain of *Xl*Bicc1, suggesting that it occurs via the canonical KH-RNA interaction mechanism used by single KH domain RNA binding proteins (Dowdle et al. 2019; Minegishi et al. 2021). The second interface required the KH1 domain but not KH1’s GXXG motif, suggesting that KH1 promoted the formation of a second GXXG-independent protein-RNA interface. Bicc1s from three additional species, human, zebrafish and fruit fly, also bound the *Xl*Cripto-1 RNA with high-affinity and specificity for the 5’ leader sequence. However, in contrast to *Xl*Bicc1, *Hs*Bicc1, *Dr*Bicc1 and *Dm*Bicc1 had a substantially relaxed requirement for the 3’ loop sequence of *Xl*Cripto-1 RNA. Together, the data support a model wherein the most conserved KH domain, KH2, forms a core, conserved Bicc1-RNA interaction, while the KH1 domain supports formation of a secondary, more diverged Bicc1-RNA interface. These findings have important implications for understanding the Bicc1-RNA interactome.

## RESULTS

### *Xl*Bicc1 bound the 32-nt Cripto-1 RNA target site with high-affinity and specificity

*Xl*Bicc1 binding to the previously defined 32-nt Cripto-1 RNA binding site was assessed using a purified recombinant protein consisting of the N-terminal region of *Xenopus laevis* Bicc1 (aa 1-506, (*Xl*Bicc1)), which encompasses the multi-KH RNA binding domain of Bicc1 (Dowdle et al., 2019) (**Fig.1A-B**) (**Fig.S1**). The *Xl*Cripto-1 RNA substrate was labeled on its 5’ end with fluorescein to allow its detection in electromobility shift (EMSA) and fluorescence polarization assays (**Fig.1B-C**) (Dowdle et al. 2017). A 32-nucleotide RNA from the 3’UTR of the *Xenopus* CyclinB1 mRNA served as the non-specific control RNA substrate (Dowdle et al. 2017). Both assays indicated high-affinity binding to the *Xl*Cripto-1 RNA target but not to the Cyclin B1 RNA by *Xl*Bicc1, as predicted from previous work (Zhang et al. 2013, 2014; Park et al. 2016; Dowdle et al. 2019). The fluorescence polarization assays allowed for determination of a K_id_ for the specific *Xl*Bicc1-*Xl*Cripto-1 complex of K_d_ = 34 nM (**Table 1**).

**Figure 1.**
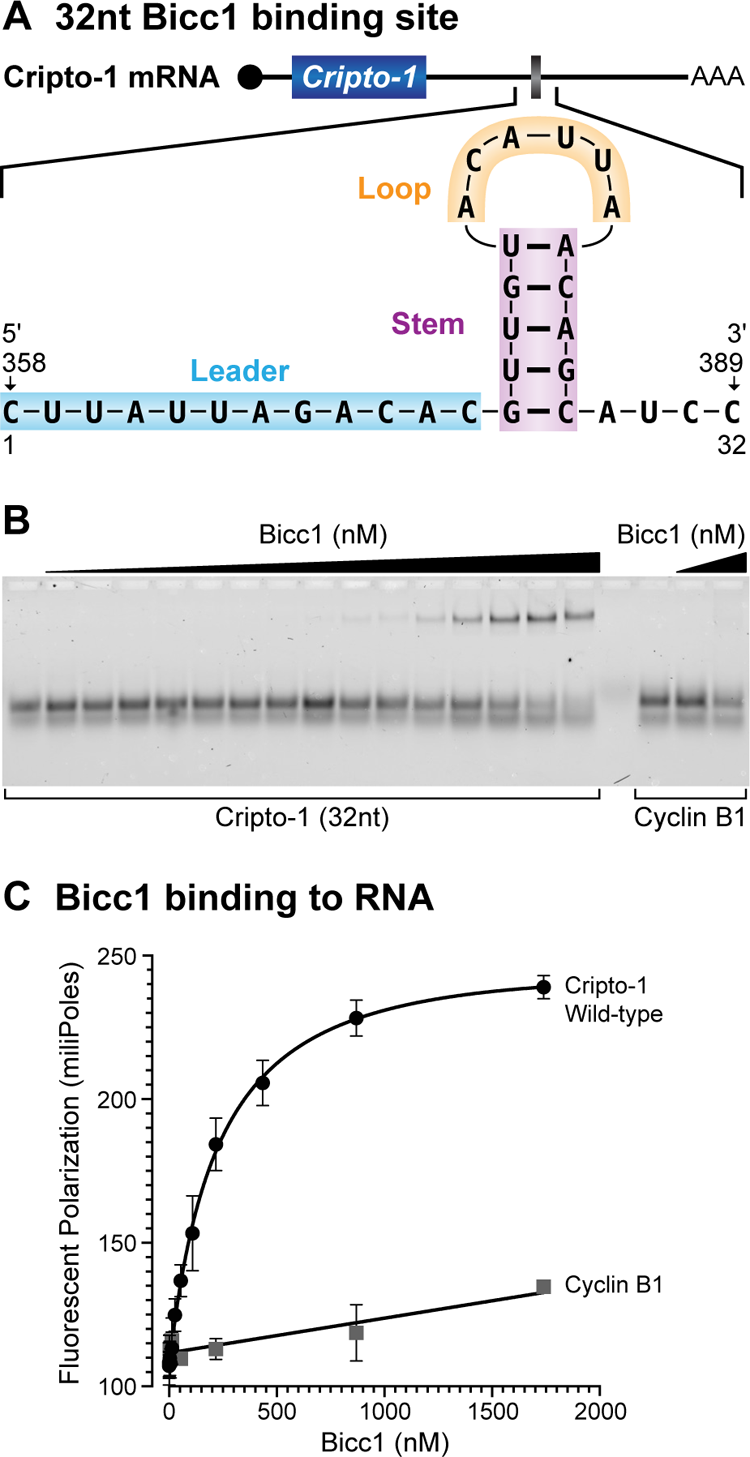
*Xenopus* Bicc1 binds with high affinity and specificity to the 32-nucleotide Bicc1 binding site within the 3’UTR of Cripto-1 mRNA. *A.* Cartoon of the *Xenopus* Cripto-1 mRNA indicating position of the minimal 32-nt Cripto-1 *Xl*Bicc1 RNA target site that functions in Bicc1-dependent translational repression (refs). This validated Bicc1 target site will be referred to as the Cripto-1 RNA site, which starts 358 nucleotides 3’ after the Cripto stop codon. The nucleotide sequence of the Cripto-1 RNA site is indicated with the leader, stem and loop regions highlighted. Previously published studies support this model for the Cripto-1 RNA site (refs). *B.* EMSA experiment was used to assess binding of *Xl*Bicc1 to a 3’-fluorescein labeled Cripto-1 RNA site. A 32-nt region from the Cyclin B1 mRNA 3’ UTR served as a negative control substrate for Bicc1 (Hereafter, referred to as Cyclin B1 negative control RNA). *C.* Fluorescent polarization was used to quantitatively assess Bicc1 to a 3’-fluorescein labeled Cripto1 RNA or Cyclin B1 negative control RNA. Data from three independent experiments were used to determine the mean value and standard deviation for each concentration of Bicc1 indicated. A K_d_ of 34 nM was determined for the XlBicc1-Cripto-1 RNA interaction (see Table 1). Cyclin B1 negative control RNA showed such weak binding that a K_d_ was unmeasurable.

**Table 1.**
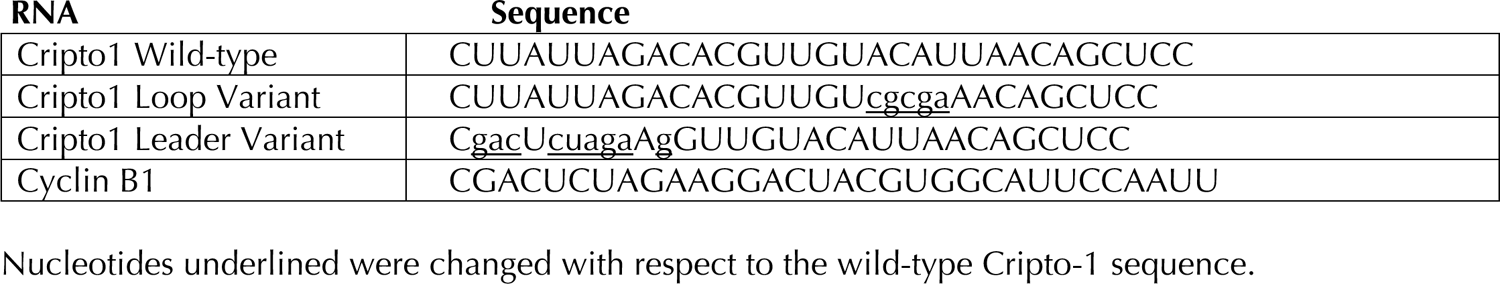
RNA substrates used in this study

### Two distinct sequence elements within the Cripto-1 RNA site were required for high-affinity binding by *Xl*Bicc1

RNA-footprinting, EMSA and translational reporter assays validate the biochemical and functional relevance of the Bicc1-binding site within the *Xl*Cripto-1 mRNA 3’ UTR (**Fig.2A**) (Zhang et al. 2014). Mutational analyses support a model for the requirement for a stem-loop structure, and indicate that the sequence of the loop is also an important determinant. However, the stem-loop does not fully explain *Xl*Bicc1 binding, suggesting additional sequence determinants are relevant (Zhang et al. 2014). Two different mutant variants of the *Xl*Cripto-1 RNA site were generated: one variant contained a complete substitution of the 3’ loop sequence while the other contained a complete substitution of the 5’ leader sequence (**Fig.2A**). Bicc1 binding to these mutant RNA substrates was examined by EMSA or fluorescence polarization (**Fig.2B-C**). The data from these assays provided evidence that both the leader and the loop sequences of the Cripto-1 RNA site were essential for high-affinity binding by *Xl*Bicc1, though the loop variant provided a minimal amount of binding energy to allow for calculation of a K_d_ by fluorescence polarization (estimated K_d_ of Bicc1 for 3’ loop mutant = 12 μM). Nevertheless, the experiments indicated that the 3’loop sequence of the *Xl*Cripto-1 RNA site was critical as the loop mutant reduced the affinity of *Xl*Bicc1 for the RNA substrate > 170-fold (K_d_ WT/ K_d_ loop mutant = 0.0057) (**Table 1**). Thus the *Xl*Bicc1 target site in the Cripto-1 RNA is, minimally, a bipartite element, with two distinct non-contiguous sequence elements contributing to the stability of the *Xl*Bicc1-Cripto-1 RNA interaction.

**Figure 2.**
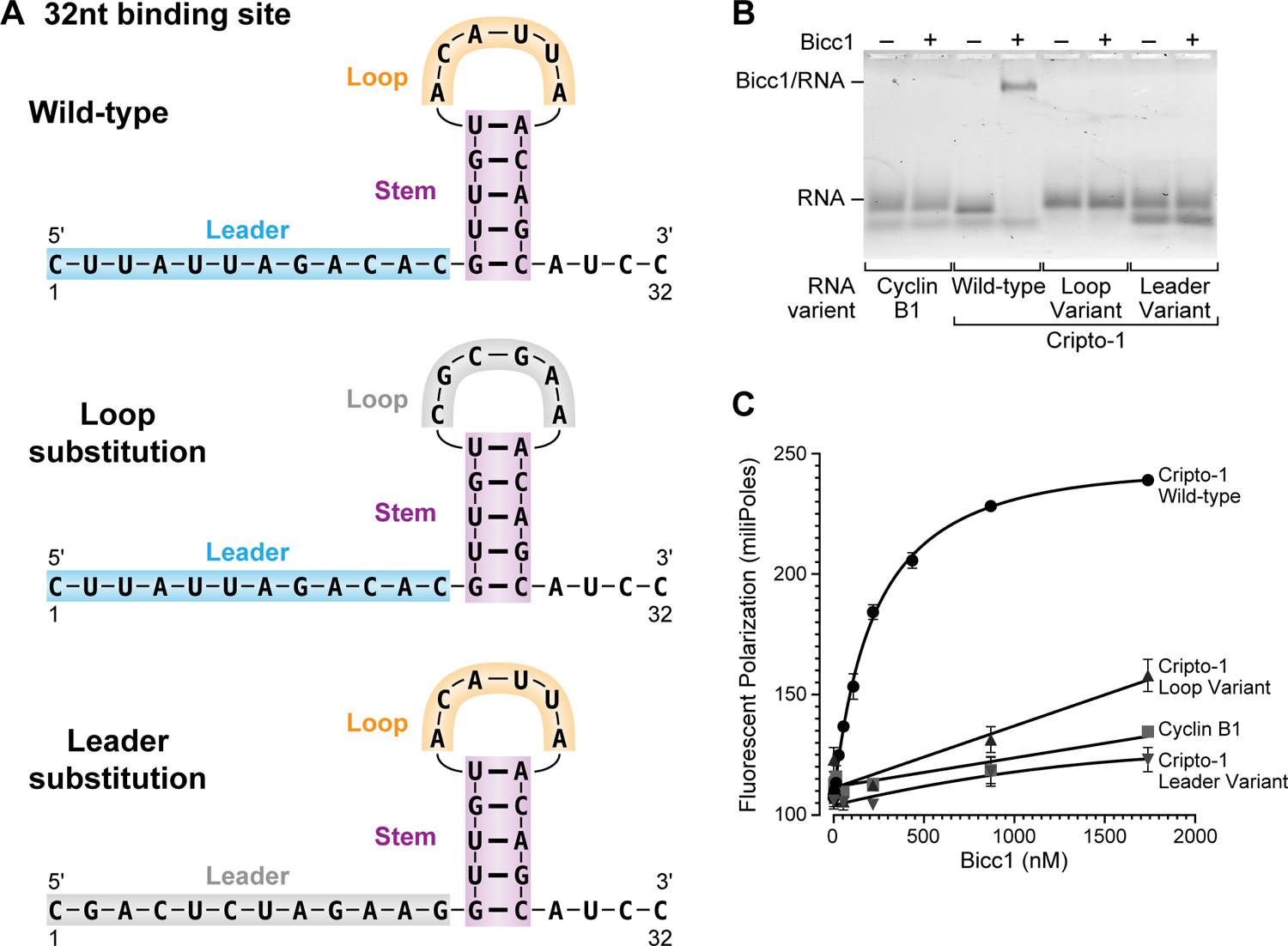
The Loop and Leader sequences were each required for high-affinity binding of the Cripto-1 RNA site by *Xl*Bicc1. ***A.*** Diagrams of the wild-type, loop-mutant (grey highlight), and leader-mutant (grey highlight) versions of the 32-nt Cripto-1 RNA *Xl*Bicc1 binding site. ***B.*** EMSA was used to assess binding of *Xl*Bicc1 to the Cripto-1 sites depicted in *A.* Cyclin B1 served as the negative control RNA. ***C.*** Fluorescent polarization was used to quantitatively assess Bicc1 to 3’-fluorescein labeled Cripto-1 RNA or Cyclin B1 negative control RNA. Data from three independent experiments were used to determine the mean value and standard deviation for each concentration of Bicc1 indicated. 350nM of purified Bicc1 protein and 10nM of fluorescein labeled RNA were used. K_d_s reported in Supplementary Table 1.

### Bicc1 from three other species bound to the *Xenopus* Cripto-1 RNA target with high-affinity

While the multi-KH domain of Bicc1 is conserved across metazoans, the KH2 domain is the most conserved subdomain at the level of amino acid identity (**Fig.3A and Fig.S1**) (Dowdle et al. 2019). In addition, the KH2 domain is the only KH domain within *Xl*Bicc1 whose conserved GXXG motif is critical for *Xl*Cripto-1 RNA binding in vitro and in vivo (Dowdle et al. 2019; Minegishi et al. 2021). Thus, only the KH2 domain interacts with the Cripto-1RNA via a canonical KH-GXXG-dependent protein-RNA interaction. The high conservation of the Bicc1 RNA binding domain, particularly over the functionally critical KH2 domain, motivated us to ask whether Bicc1’s derived from three other species, human (*Hs*Bicc1), zebrafish (*Dr*Bicc1) and fruit fly (*Dm*Bicc1), could bind to the *Xl*Cripto-1 RNA site (**Fig.3B**). Purified recombinant Bicc1 proteins were used in fluorescence polarization assays as described above (**Fig.2., Fig.3B**). EMSA assays were also performed as an independent assessment (**Fig.S3**). Both RNA binding assays revealed that Bicc1 derived from human, zebrafish and fruit fly bound to the *Xl*Cripto-1 RNA with high affinity and specificity compared to the Cyclin B1 negative binding control RNA (**Fig.3B-C, Table 1**). These data provided evidence that an important feature(s) of the Bicc1-RNA substrate interface was conserved among metazoans.

**Figure 3.**
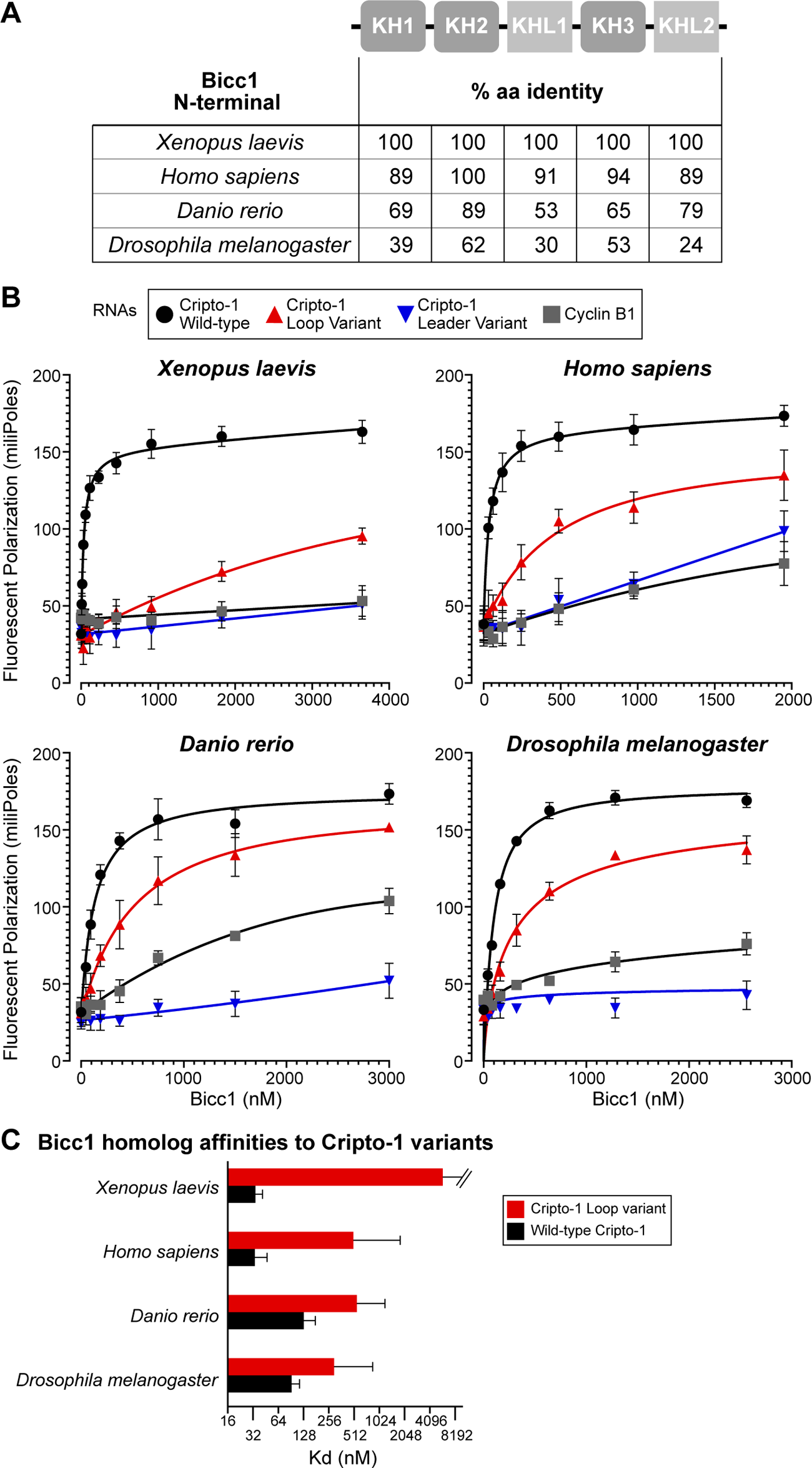
Specific and high-affinity binding of the defined 32-nt Cripto-1 RNA binding site for *Xl*Bicc1 by HsBicc1, DrBicc1, DmBicc1. ***A.*** Percentage of amino acid identity for the indicated KH1 and KHL domains within the defined Bicc1 RNA binding module between XlBicc1 and Bicc1s from the indicated species. Clustal Omega was used for these analyses. ***B.*** Fluorescent polarization was used to assessed binding of the indicated specie’s Bicc1 to the 3’-fluorescein labeled wild-type and mutant versions of *Xl*Cripto-1 RNA sites as described in Figure 2A. Data from three independent experiments were used to determine the mean value and standard deviation for each concentration of Bicc1 indicated. K_d_ s and additional relevant values reported in Table 1. ***C.*** A bar graph comparing the measured K_d_s for the Bicc1 homologs affinities for the Wild-type *Xl*Cripto-1 (black) and *Xl*Cripto-1 loop variant (red) RNAs. The K_d_s were measured from at least three independent experiments with error bars representing the 95% confidence interval. K_d_ values are reported in Supplementary table 1.

### The non-*Xenopus* Bicc1 proteins bound the *Xl*Cripto-1 RNA with specificity for the 5’ leader sequence but a relaxed requirement for the 3’ loop sequence

To ask whether non-*Xenopus* Bicc1 proteins interacted with the *Xl*Cripto-1 RNA substrate via conserved *Xl*Bicc1-RNA interfaces, the 5’ leader and 3’ loop Cripto-1 RNA variants were assessed for binding by *Hs*Bicc1, *Dr*Bicc1 and *Dm*Bicc1(**Fig.3B-C**). As seen for *Xl*Bicc1, the 5’ leader variant Cripto-1 RNA abolished Bicc1-RNA complex formation with each of the non-*Xenopus* Bicc1 proteins. Thus, the 5’ leader sequence was required for high-affinity binding with all the species’ Bicc1s, providing evidence that it defined a core, conserved RNA interface for Bicc1. However, while the 3’ loop variant Cripto-1 RNA reduced Bicc1-RNA complex formation with each of the non-*Xenopus* Bicc1 proteins, the negative effects on Bicc1-RNA stability were mild compared to the effects this same variant had on *Xl*Bicc1 binding (**Fig.3B-C**). Thus, each of the non-*Xenopus* Bicc1 proteins had a relaxed requirement for the 3’ loop sequence relative to *Xl*Bicc1.

Specifically, the 3’ loop variant *Xl*Cripto-1 RNA reduced the binding strength of *Xl*Bicc1 by >170-fold relative to the wild-type *Xl*Cripto-1 RNA (**Fig.2C and Table 1**). In contrast, the 3’ loop variant of *Xl*Cripto-1 RNA reduced the binding strength of the *Hs*Bicc1, *Dr*Bicc1, and *Dm*Biccl1 only minimally relative to the wild-type Cripto-1 RNA, by 15-fold, 4.3-fold and 3.2 fold, respectively (**Fig.3C and Table 1**). Thus the Bicc1-RNA interaction that involved the 5’ leader sequence was a substantially more conserved contributor to Bicc1-RNA binding strength across species compared to the Bicc1-RNA interaction that involved the 3’ loop sequence.

### The KH1 domain of *Xl*Bicc1 promoted the strong requirement of *Xl*Bicc1 for the 3’ loop region of *Xl*Cripto-1 RNA

The RNA binding experiments with the non-*Xenopus* Bicc1 proteins indicated that *Xl*Bicc1 was distinctly capable of promoting a robust Bicc1-RNA interaction with the 3’ loop region of *Xl*Cripto-1 RNA that contributed to the stability of the *Xl*Bicc1-*Xl*Cripto-1 RNA complex. In previous work, the strong conservation of the KH2 domain helped guide us to understanding why the conserved GXXG motif within the KH2 domain was the only GXXG within Bicc1 essential for binding to *Xl*Cripto-1 RNA (Dowdle et al. 2019; Minegishi et al. 2021). Here, we again exploited clues from differences in the levels of conservation among individual sub-domains within Bicc1’s complex RNA binding domain, which led us to focus on the KH1 domain. Specifically, protein alignments revealed that in contrast to the KH2 domain, the KH1 domains from non-*Xenopus* Bicc1 proteins were divergent at the level of amino acid identity (**Fig.3A and Fig.S1**). The effect was particularly striking for *Drosophila* Bicc1. Whereas the *Xenopus* and *Drosophila* KH2 domains share 62% amino acid identity, their KH1 domains share only 39% identity (**Fig.3A and Fig.S1**). *Dm*Bicc1 also demonstrated the least dependence for the loop sequence for binding to *Xl*Cripto-1 RNA, with the loop variant reducing *Dm*Bicc1-RNA binding strength by only ∼3-fold (compared to a ∼170-fold reduction caused by this same mutant in *Xl*Bicc1-RNA binding strength) (**Fig.3B-C, Table 1**).

To examine the contribution of the KH1 domain to Bicc1 RNA binding specificity, RNA binding assays were performed with two different mutant *Xenopus* Bicc1 proteins: one mutant lacked the KH1 domain entirely (*Xl*Bicc1-ΔKH1) and the other contained an inactivating substitution of the GXXG motif, (GKSGàGDDG; this mutant referred to as *Xl*Bicc1-KH1-GDDG) (**Fig.4A**). Deletion of the KH1 domain reduced the ability of *Xl*Bicc1 to bind to the wild-type Cripto-1 RNA by 22-fold (**Fig.4B-D, Table 2**). Thus an intact KH1 domain was required for full binding energy of *Xl*Bicc1-*Xl*Cripto-1 RNA complex, as expected (Dowdle et al. 2019). In addition, *Xl*Bicc1-ΔKH1-Cripto-1 RNA binding strength remained fully dependent on the 5’ leader sequence as the relevant 5’ leader variant RNA abolished binding (**Fig.4B, blue curve**). However, *Xl*Bicc1-ΔKH1 bound to the 3’ loop mutant with only a 2-fold reduction in binding strength (**Fig.4B red curve, Fig.4D and Table 2**). Thus the 3’ loop sequence of *Xl*Cripto-1 RNA contributed minimally to the binding energy of the *Xl*Bicc1-ΔKH1-Cripto-RNA complex, providing evidence that the protein-RNA interface between *Xl*Bicc1 and the 3’ loop sequence of Cripto-1 RNA required the KH1 domain. The paradigmatic protein-RNA interface used by single KH domain proteins is the GXXG motif, which is GKSG in *Xl*Bicc1’s KH1 domain (**Fig.S1**). To test whether KH1’s role in recognition of the 3’ loop within Cripto-1 RNA was GXXG-dependent, RNA binding by the *Xl*Bicc1-KH1-GDDG mutant was also assessed (**Fig.4C-D, Table 2**). In contrast to the 22-fold reduction in binding strength to the wild-type Cripto-1 RNA caused by the *Xl*Bicc1-ΔKH1 mutant, the *Xl*Bicc1-KH1-GDDG mutant only reduced the binding strength by 1.6-fold (**Table 2**). However, the stability of the *Xl*Bicc1-KH1-GDDG-Cripto-1 RNA complex was now fully dependent on the 3’ loop sequence, as binding between the KH1-GDDG mutant protein and the 3’ loop variant was so weak that a K_d_ could not be determined (**Table 2**). Thus, the KH1 domain was required for *Xl*Bicc1’s recognition of the 3’ loop sequence with the *Xl*Cripto-1 RNA, while being largely dispensable for *Xl*Bicc1’s recognition of the 5’ leader sequence.

**Figure 4.**
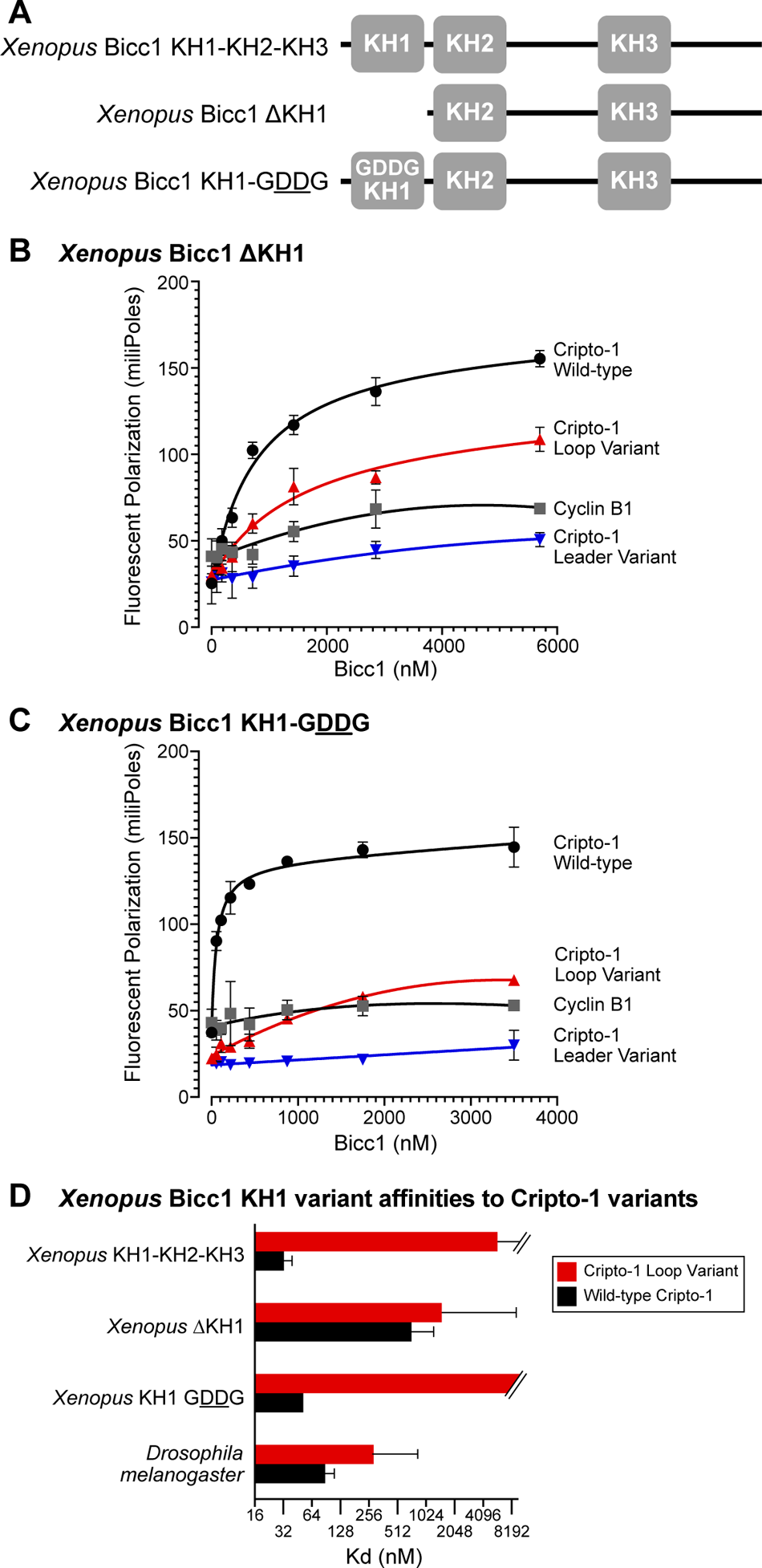
The KH1 domain, but not its canonical GXXG motif, of *Xl*Bicc1 contributed to *Xl*Bicc1’s reliance on the loop sequence for high affinity binding to the Cripto-1 RNA site. *A.* Wild-type and mutant *Xl*Bicc1 proteins used to assess binding to *Xl*Cripto-1 RNAs in Figure 2A. Fluorescent polarization was used to assess binding of the: *B. Xl*Bicc1 ΔKH1 and *C.* XlBicc1 KH1-GDDG proteins to indicated 3’-fluorescein labeled RNA substrates. Data from at least three independent experiments were used to determine the mean value and standard deviation for each concentration of Bicc1 indicated. K_d_s and additional relevant values reported in Table 2. *D.* A bar graph comparing the measured K_d_s for the for the *Xl*Bicc1 ΔKH1, XlBicc1 KH1-GDDG compared to the Wild type *Xl*Bicc1 Kh1-Kh2-K3 protein for the Wild-type *Xl*Cripto-1 (black) and *Xl*Cripto-1 loop variant (red) RNAs. The K_d_s were measured from at least three independent experiments with error bars representing the 95% confidence interval. K_d_ values are reported in Supplementary Table 2.

### The KH1 domain of *Xl*Bicc1 was sufficient to confer 3’ loop-dependent binding to *Dm*Bicc1

The *Dm*Bicc1 protein interacted with the *Xl*Cripto-1 RNA similarly to how the *Xl*Bicc1-τ−KH1 mutant did, suggesting that something about the *Dm*Bicc1’s KH1 domain prevented it from recognizing the 3’ loop. To test this idea, we generated recombinant fusion proteins between *Xl*Bicc1 and *Dm*Bicc1 and asked how these engineered proteins interacted with *Xl*Cripto-1 RNA substrates (**Fig.5A**). Importantly, *Dm*Bicc1 and *Xl*KH1-*Dm*Bicc1KH2-KH3, an engineered protein fusion where the N-terminal region encoding the *Dm*Bicc1-KH1 domain was substituted with the corresponding *Xl*Bicc1 region, bound to the wild-type Cripto-1 RNA with indistinguishable binding strengths (*Dm*Bicc1 K_d_ = 93nM and *Xl*KH1-DmBicc1KH2-KH3 K_d_ = 89nM) (**Fig.5B and E, Table 3**). However, while *Dm*Bicc1 was only minimally dependent on the 3’ loop sequence for stable binding, the *Xl*KH1-*Dm*Bicc1KH2-KH3 fusion protein was fully dependent on this sequence for detectable binding (**Fig.5B and E, Table 3**). A caveat to interpreting this experiment was that the *Xl*KH1-*Dm*KH1 domain swap included a short intrinsically disordered region (IDR) N-terminal to the KH1 domain that is substantially diverged between different species’ Bicc1s. Therefore, it was possible that the IDR, and not the KH1 domain, was primarily responsible for the shift in the fusion protein’s dependence on the 3’ loop sequence for *Xl*Cripto-1 RNA binding. However, a swap between the *Xl*IDR and *Dm*IDR produced a *Xl*IDR-*Dm*KH1-KH3 fusion protein that behaved indistinguishably from *Dm*Bicc1 in terms of binding strengths for the wild-type and variant *Xl*Cripto-1 RNA substrates (**Fig.5A, C, E, Table 3**). Conversely, the opposite swap of these organism’s Bicc1 IDRs produced a *Dm*IDR-*Xl*KH1-KH3 fusion protein that behaved similarly to *Xl*Bicc1, showing a substantial dependence on the 3’ loop sequence of *Xl*Cripto-1 RNA for stable *Xl*Bicc1-*Xl*Cripto-1 RNA complex formation (**Fig.5A, D-E, Table 3**). Taken together, these data provided evidence that the *Xl*KH1 domain was both necessary and sufficient, in the context of *Dm*Bicc1, to confer Bicc1’s dependence on the 3’ loop sequence of the *Xl*Cripto-1-RNA for stable Bicc1-RNA binding (**Fig.5E, Table 3**). Thus the *Xl*Bicc1 KH1 domain promoted a Bicc1-RNA interface dependent on the 3’ loop sequence of the *Xl*Cripto-1 RNA *Xl*Bicc1 target site.

**Figure 5.**
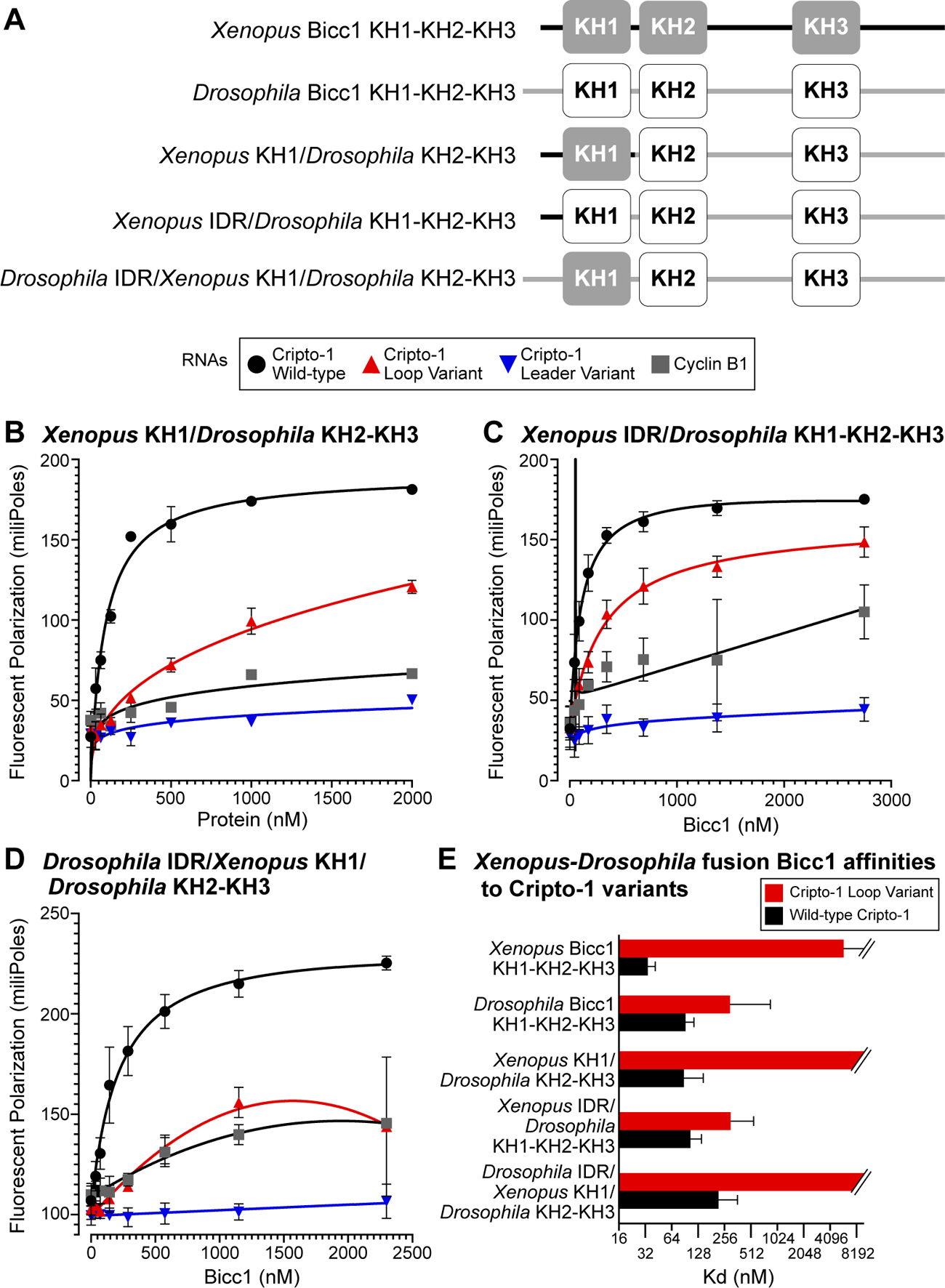
The KH1 domain of *Xl*Bicc1 is sufficient to confer a requirement for the XlCripto-1 RNA loop sequence to DmBicc1 KH2-KH3. ***A.*** *Xl*Bicc/*Dm*Bicc1 variants used in these experiments. Fluorescent polarization was used to assess binding of the: ***B.*** *Xl*KH1-*Dm*KH2-KHL2, ***C,*** *Xl*IDR-*Dm*KH1-KHL2, and ***D.*** *Dm*IDR-*Xl*KH1-KHL2 variant proteins. Data from at least six independent experiments were used to determine the mean value and standard deviation for each concentration of Bicc1 indicated. K_d_s and additional relevant values reported in Table 3. ***E.*** A bar graph comparing the measured K_d_s for the *Xenopus-Drosophila* Bicc1 fusion proteins for the wild-type *Xl* Cripto-1 (black) and *Xl* Cripto-1 loop variant (red) RNAs. The K_d_s were measured from at least three independent experiments with error bars representing the 95% confidence interval. K_d_ values are reported in Supplementary Table 3.

## DISCUSSION

Bicc1 is conserved protein that plays a critical role in establishing the anterior-posterior body plan during metazoan embryogenesis (Gamberi and Lasko 2012; Mahone et al. 1995; Wessely and De Robertis 2000; Wessely et al. 2001; Park et al. 2016). Bicc1 also functions in adult organisms to control tissue homeostasis, with Bicc1 mutations linked to polycystic kidney disease, an aggressive form of liver cancer and major depressive disorder (MDD), a severe form of depression (Cogswell et al. 2003; Bouvrette et al. 2008, 2010; Tran et al. 2010; Rothé et al. 2020; Arai et al. 2014; Ying et al. 2019; Cristinziano et al. 2021; Lewis et al. 2010; Bermingham et al. 2012; Ota et al. 2015; Davidson et al. 2016; Ryan et al. 2016; Wang et al. 2019). It is clear that Bicc1’s biological roles require that it functions as an RNA binding protein to select specific target mRNAs for translational repression. A strong example is the Bicc1-mediated repression of Cripto-1 mRNA translation during the maternal stages of embryogenesis in *Xenopus laevis* that we have studied for many years (Zhang et al. 2013, 2014; Park et al. 2016; Dowdle et al. 2019). Specifically, Bicc1 dictates the anterior-posterior body plan in vertebrates in large part because it binds to a target site within the 3’ UTR of Cripto-1 mRNA and inhibits Cripto-1 mRNA translation in specific cells of the developing embryo. This Bicc1-mediated repression thereby restricts Cripto-1 protein accumulation to only those cells destined to become the future ectoderm and anterior cells of the organism. Thus Bicc1’s ability to select specific target RNAs for translational repression defines its own role a cell-fate regulator. However, despite Bicc1’s biological importance and the salient role of its selective RNA binding function, fundamental information about how Bicc1 recognizes target mRNAs is lacking. A challenge is that only a small number of Bicc1 target sites have been reported, and even fewer have been biologically validated or characterized in sufficient detail to be most informative in quantitative Bicc1-RNA interaction studies. We exploited the minimal *Xl*Bicc1 target site within the *Xl*Cripto-1 mRNA that we have identified and extensively validated functionally both in vitro and in vivo (Zhang et al. 2014) and **Fig.1A**). This Bicc1-target site is found within the 3’ UTR of the *Xl*Cripto-1 mRNA, and consists of 32 contiguous nucleotides comprised of a 5’ leader sequence followed by a 3’ stem-loop structure, where the structure of the stem, and not the sequence is important (Zhang et al. 2014). Fluorescence polarization was used to quantify the binding of recombinant, purified Bicc1 proteins derived from four species, *Xenopus laevis*, *Homo sapien*, *Danio rerio* and *Drosophila melanogaster* to wild-type and two engineered variant *Xl*Cripto-1 RNA targets, a 5’ leader sequence variant and 3’ loop sequence variant. The data supported four key conclusions. First, *Xl*Bicc1 binds to its native *Xl*Cripto-1 target site with a high-affinity and specificity involving at least two distinct regions of the RNA substrate, the 5’ leader and 3’ loop sequences. Second, the 5’ leader sequence defines an evolutionarily conserved RNA interface because each of the other species Bicc1s could bind the *Xl*Cripto-1 RNA target site with a high-affinity that required the 5’ leader sequence. Third, and in contrast to the 5’ leader sequence requirement, the 3’ loop sequence provided only minimal binding energy to the non-*Xenopus*-Bicc1-RNA complexes. These data provided evidence that the *Xl*Cripto-1 RNA binding site functioned as a bipartite element that provided for the formation of two distinct protein-RNA interfaces involving Bicc1. Fourth, the most diverged of the subdomains that comprise the multi-KH and KHL RNA binding domain of Bicc1, KH1, was key for endowing *Xl*Bicc1 with its strong dependence on the 3’ loop sequence. Together with our previous analyses of *Xl*Bicc1-RNA interactions, we propose that selective, high-affinity Bicc1-RNA binding requires two distinct protein-RNA interfaces, one requiring the highly conserved KH2 domain and the 5’ leader sequence that occurs via the canonical GXXG-dependent mechanism, and a second, more diverged Bicc1-RNA interface, that requires the 3’ loop sequence. While information about direct protein-RNA contacts require further investigation, the data presented in this report support the conclusion that the KH1 subdomain is important for providing for a secondary, more evolutionarily flexible Bicc1-RNA interface. This secondary interface varies between species, but may also vary considerably between RNA substrates to help fine-tune the breadth and strength of Bicc1-mediated translational repression within a cell and in turn fine tune the composition of the cell’s proteome.

The conclusion that the Bicc1 binding site within the *Xl*Cripto-1 RNA acts as a bipartite element requiring a sequence-independent stem structure is consistent with a recent large-scale biochemical study of mammalian RNA binding proteins that revealed that several multi-KH domain proteins require relatively large RNA recognition sites that are often bipartite and include stem-loops (Lewis et al. 2000; Du et al. 2007; Chao et al. 2010; Nicastro et al. 2017; Yang et al. 2017; Dominguez et al. 2018; Biswas et al. 2019). In addition, another recent study concludes that mouse Bicc1 recognizes a target RNA, Dand5, in a mechanism that requires two tandem GAC motifs, consistent with the existence of a bipartite target site. Interestingly, the 5’ leader sequence of the *Xl*Cripto-1 RNA target site contains a GAC motif. However, this short motif can also be found in the negative control RNA used in this study, Cyclin B1, and that the 5’ leader sequence of Cripto-1 RNA is insufficient on its own to bind *Xl*Bicc1 (Zhang et al. 2014). Finally, a structural study of a Bicc1-related multi-KH-domain protein from *C. elegans* reveals that the region of this protein analogous to the KH2-KHL1-KH3-KHL2 region of Bicc1 forms a single, compact domain, while the analogous KH1 domain forms a distinct structural appendage (Nakel et al. 2010). Thus, while further investigations are required, together with the data presented in this report, recent studies are suggesting the emergence of generalizable features of Bicc1-RNA interactions that will help clarify and define the important Bicc1-RNA interactome.

## METHODS AND MATERIALS

### Bicc1 protein expression and purification

Bicc1 proteins were cloned into pET28b Bacterial expression vectors as N-terminal fusions with a His-6, SUMO tag (Malakhov et al. 2004). Cultures of *E.coli* cells containing each plasmid were grown to an OD600 of 0.6 and induced with 1mM IPTG at 25 degrees C overnight. The cells were collected and lysed in B-PER reagent, 1/2X TBS, 2.5mM MgCl2, 2.5% glycerol, 1mM BME, 10mM Imidazole, 200mM NaCl, 1mM ATP, and protease inhibitors. The soluble lysate was applied to a HisTrap FF chromatography column. The column was washed 10 times with 1xTBS, 5mM MgCl2, 5% glycerol, 2mM BME, 20mM Imidazole, 400mM NaCl and the proteins were then eluted with 450mM imidazole and dialyzed in 1x TBS.

### Proteins

The sequences of the various purified Bicc1 proteins used are indicated below. The SUMO and 6-His tags are highlighted in lowercase.

### SUMO-*Xl*-Bicc1 Nterm (KH1-KH3)

msdsevnqeakpevkpevkpethinlkvsdgsseiffkikkttplrrlmeafakrqgkemdslrflydgiriqadqtpedldmedndiieahre qiggenlyfqAAQCESIGGDMNQSDPGSNSERSADSPVPGSEDDSPHDPEWREERFRVDRKKLETMLQAAA EGKGKSGEDFFQKIMEETNTQIAWPSKLKIGAKSKKDPHIKVSGKKENVKEAKERIMSVLDTKSNRVTLKMDVSHTEHSHVIGKGGNNIKKVMEETGCHIHFPDSNRNNQAEKSNQVSIAGQPAGVESARVRIRELLPLVLMFELPIAGILQPIPDPNSPTIQQISQTYNLTVSFKQRSRVYGATVIVRGSQNNTSAVKEGTAMLLEHLAGSLATAIPVSTQLDIAAQHHLFMMGRNGCNIKHIMQRTGAQIHFPDPNNPLKKSTVYLQGTIDSVCLARQYLMGCLPLVLMFDMKEEIEIEPQCITQLMEQLDVFISIKPKPKQPSKSVIVKSVERNALNMYEARKCLLGLDSSGVTVTNQSTLSCPVPMNCHGLDILAAGLGLSGLGLLGPNTLSVNSTTAQNSLLNALNSSLSPLHSPSSAAPS PTLWAASLANNATGFENLYFQAhhhhhh

### SUMO-*Xl*-Bicc1 Nterm (KH1GDDG KH2-KH3)

msdsevnqeakpevkpevkpethinlkvsdgsseiffkikkttplrrlmeafakrqgkemdslrflydgiriqadqtpedldmedndiieahre qiggenlyfqAAQCESIGGDMNQSDPGSNSERSADSPVPGSEDDSPHDPEWREERFRVDRKKLETMLQAAA EGKGKSGEDFFQKIMEETNTQIAWPSKLKIGAKSKKDPHIKVSGKKENVKEAKERIMSVLDTKSNRVTLKMDVSHTEHSHVIGDDGNNIKKVMEETGCHIHFPDSNRNNQAEKSNQVSIAGQPAGVESARVRIRELLPLVLMFELPIAGILQPIPDPNSPTIQQISQTYNLTVSFKQRSRVYGATVIVRGSQNNTSAVKEGTAMLLEHLAGSLATAIPVSTQLDIAAQHHLFMMGRNGCNIKHIMQRTGAQIHFPDPNNPLKKSTVYLQGTIDSVCLARQYLMGCLPLVLMFDMKEEIEIEPQCITQLMEQLDVFISIKPKPKQPSKSVIVKSVERNALNMYEARKCLLGLDSSGVTVTNQSTLSCPVPMNCHGLDILAAGLGLSGLGLLGPNTLSVNSTTAQNSLLNALNSSLSPLHSPSSAAP SPTLWAASLANNATGFENLYFQAhhhhhh

### SUMO-*Xl*-Bicc1 Nterm (KH2-KH3)

msdsevnqeakpevkpevkpethinlkvsdgsseiffkikkttplrrlmeafakrqgkemdslrflydgiriqadqtpedldmedndiieahre qiggenlyfqSHTEHSHVIGDDGNNIKKVMEETGCHIHFPDSNRNNQAEKSNQVSIAGQPAGVESARVRIRE LLPLVLMFELPIAGILQPIPDPNSPTIQQISQTYNLTVSFKQRSRVYGATVIVRGSQNNTSAVKEGTAMLLEHLAGSLATAIPVSTQLDIAAQHHLFMMGRNGCNIKHIMQRTGAQIHFPDPNNPLKKSTVYLQGTIDSVCLARQYLMGCLPLVLMFDMKEEIEIEPQCITQLMEQLDVFISIKPKPKQPSKSVIVKSVERNALNMYEARKCLLGLDSSGVTVTNQSTLSCPVPMNCHGLDILAAGLGLSGLGLLGPNTLSVNSTTAQNSLLNALNSSLSPLHS PSSAAPSPTLWAASLANNATGFENLYFQAhhhhhh

### SUMO-*Hs*-Bicc1 Nterm

msdsevnqeakpevkpevkpethinlkvsdgsseiffkikkttplrrlmeafakrqgkemdslrflydgiriqadqtpedldmedndiieahre qiggenlyfqAAQGEPGYLAAQSDPGSNSERSTDSPVPGSEDDLVAGATLHSPEWSEERFRVDRKKLEAMLQ AAAEGKGRSGEDFFQKIMEETNTQIAWPSKLKIGAKSKKDPHIKVSGKKEDVKEAKEMIMSVLDTKSNRVTLKMDVSHTEHSHVIGKGGNNIKKVMEETGCHIHFPDSNRNNQAEKSNQVSIAGQPAGVESARVRIRELLPLVLMFELPIAGILQPVPDPNSPSIQHISQTYNISVSFKQRSRMYGATVIVRGSQNNTSAVKEGTAMLLEHL AGSLASAIPVSTQLDIAAQHHLFMMGRNGSNIKHIMQRTGAQIHFPDPSNPQKKSTVYLQGTIESVCLARQYLMGCLPLVLMFDMKEEIEVDPQFIAQLMEQLDVFISIKPKPKQPSKSVIVKSVERNALNMYEARKCLLGLESSGVTIATSPSPASCPAGLACPSLDILASAGLGLTGLGLLGPTTLSLNTSTTPNSLLNALNSSVSPLQENLYF QAhhhhhh

### SUMO-*Dr*-Bicc1 Nterm

msdsevnqeakpevkpevkpethinlkvsdgsseiffkikkttplrrlmeafakrqgkemdslrflydgiriqadqtpedldmedndiieahre qiggenlyfqAAEPLSFMHHDHGSSSERSDDSPSAVSEDDSSGHCGHISPPDPDWTEERFRVDRKKLETMLLA ANEGRINGDDFFQKVMDETNTQIAWPSKLKIGAKSKKDPHIKVSGKRDDVREAKEKIMSVLDTKSHRVTLKMDVSHTEHSHVIGKGGHNIKRVMEETGCHIHFPDSNRHSQAEKSNQVSIAGQLTGVEAARVKIRELLPLVLMFECSGVVQLVDCSSPVVQHISHTYNVSISFRPPSRLYGNTAIVRANQNNSSGVKRGTALLLEHLAGSLASSVMVSTQLDIAPQHHHFLLGRNGANIKLISQRTGAHIHFPEISPHNSNASRSAVYIQGSIDAVCAARQQIMGCLPLVLLFDIKEETEVASQVITTLMEQLDVFISIKPKPKQPSKSVIVKSVERNAGSLYEVRRILLGLESSCLSSSVSSVSVNGHISSSPPRIASIGLDTLASAGLRLSTLADLLRSSVSPVPNGSPNPSCALNGHGSVLNIQNGV NNTQHIHENLYFQAhhhhhh

### SUMO-*Dm*-Bicc1 Nterm

msdsevnqeakpevkpevkpethinlkvsdgsseiffkikkttplrrlmeafakrqgkemdslrflydgiriqadqtpedldmedndiieahre qiggenlyfqAMLSCASFNKLMYPSAADVAKPPMVGLEVEAGSIGSLSSLHALPSTTSVGSGAPSETQSEISSVD SDWSDIRAIAMKLGVQNPDDLHTERFKVDRQKLEQLIKAESSIEGMNGAEYFFHDIMNTTDTYVSWPCRLKIGAKSKKDPHVRIVGKVDQVQRAKERILSSLDSRGTRVIMKMDVSYTDHSYIIGRGGNNIKRIMDDTHTHIHFPDSNRSNPTEKSNQVSLCGSLEGVERARALVRLSTPLLISFEMPVMGPNKPQPDHETPYIKMIETKFNVQVIFSTRPKLHTSLVLVKGSEKESAQVRDATQLLINFACESIASQILVNVQMEISPQHHEIVKGKNNVNLLSIMERTQTKIIFPDLSDMNVKPLKKSQVTISGRIDDVYLARQQLLGNLPVALIFDFPDNHNDASEIMSLNT KYGVYITLRQKQRQSTLAIVVKGVEKFIDKIYEARQEILRLATPFVKPEIPDYYFMPKDKDLNLAYRTQLTAL LAGYVDSENLYFQAhhhhhh

### SUMO-*Xl* (aa1-134 KH1)-*Dm*KH2KH3

msdsevnqeakpevkpevkpethinlkvsdgsseiffkikkttplrrlmeafakrqgkemdslrflydgiriqadqtpedldmedndiieahre qiggenlyfqAMAAQCESIGGDMNQSDPGSNSERSADSPVPGSEDDSPHDPEWREERFRVDRKKLETMLQA AAEGKGKSGEDFFQKIMEETNTQIAWPSKLKIGAKSKKDPHIKVSGKKENVKEAKERIMSVLDTKSNRVIMKMDVSYTDHSYIIGRGGNNIKRIMDDTHTHIHFPDSNRSNPTEKSNQVSLCGSLEGVERARALVRLSTPLLISFEMPVMGPNKPQPDHETPYIKMIETKFNVQVIFSTRPKLHTSLVLVKGSEKESAQVRDATQLLINFACESIASQILVNVQMEISPQHHEIVKGKNNVNLLSIMERTQTKIIFPDLSDMNVKPLKKSQVTISGRIDDVYLARQQLLGNLPVALIFDFPDNHNDASEIMSLNTKYGVYITLRQKQRQSTLAIVVKGVEKFIDKIYEARQEILRLATP FVKPEIPDYYFMPKDKDLNLAYRTQLTALLAGYVDSENLYFQAhhhhhh

### SUMO-*Xl*IDR-*Dm*KH1KH3

msdsevnqeakpevkpevkpethinlkvsdgsseiffkikkttplrrlmeafakrqgkemdslrflydgiriqadqtpedldmedndiieahre qiggenlyfqAMAAQCESIGGDMNQSDPGSNSERSADSPVPGSEDDSPHDPEWREERFKVDRQKLEQLIKA ESSIEGMNGAEYFFHDIMNTTDTYVSWPCRLKIGAKSKKDPHVRIVGKVDQVQRAKERILSSLDSRGTRVIMKMDVSYTDHSYIIGRGGNNIKRIMDDTHTHIHFPDSNRSNPTEKSNQVSLCGSLEGVERARALVRLSTPLLISFEMPVMGPNKPQPDHETPYIKMIETKFNVQVIFSTRPKLHTSLVLVKGSEKESAQVRDATQLLINFACESIASQILVNVQMEISPQHHEIVKGKNNVNLLSIMERTQTKIIFPDLSDMNVKPLKKSQVTISGRIDDVYLARQQLLGNLPVALIFDFPDNHNDASEIMSLNTKYGVYITLRQKQRQSTLAIVVKGVEKFIDKIYEARQEILRLA TPFVKPEIPDYYFMPKDKDLNLAYRTQLTALLAGYVDSENLYFQAhhhhhh

### SUMO-*Dm*IDR-*Xl*KH1-*Dm*KH2KH3

msdsevnqeakpevkpevkpethinlkvsdgsseiffkikkttplrrlmeafakrqgkemdslrflydgiriqadqtpedldmedndiieahre qiggenlyfqAMLSCASFNKLMYPSAADVAKPPMVGLEVEAGSIGSLSSLHALPSTTSVGSGAPSETQSEISSVD SDWSDIRAIAMKLGVQNPDDLHTERFRVDRKKLETMLQAAAEGKGKSGEDFFQKIMEETNTQIAWPSKLKIGAKSKKDPHIKVSGKKENVKEAKERIMSVLDTKSNRVIMKMDVSYTDHSYIIGRGGNNIKRIMDDTHTHIHFPDSNRSNPTEKSNQVSLCGSLEGVERARALVRLSTPLLISFEMPVMGPNKPQPDHETPYIKMIETKFNVQVIFSTRPKLHTSLVLVKGSEKESAQVRDATQLLINFACESIASQILVNVQMEISPQHHEIVKGKNNVNLLSIMERTQTKIIFPDLSDMNVKPLKKSQVTISGRIDDVYLARQQLLGNLPVALIFDFPDNHNDASEIMSLNTKYGVYITLRQKQRQSTLAIVVKGVEKFIDKIYEARQEILRLATPFVKPEIPDYYFMPKDKDLNLAYRTQLTALL AGYVDSENLYFQAhhhhhh

### EMSA

Recombinant SUMO-Bicc1 N-terminal proteins were expressed and purified as described above. The 32-nucleotide Cripto1 and CyclinB1 3’-fluorescein-labeled RNA substrates were purchased from IDT (**Table 4**). Binding reactions (50μl) contained SUMO-Bicc1 protein, 10mM Heps, pH 7.5, 1mM EDTA, 50 mM KCl, 0.02% Tween 20, 0.1mg/ml yeast tRNA, 100μg/ml BSA, 2mM DTT, and 10nM RNA (Dowdle et al 2019. Reaction products were analyzed on 7.5% (1× TBE) native polyacrylamide gels (Dowdle et al. 2017). The gels were then scanned at 473 nm using a fluorimager.

### Fluorescent Polarization assays for RNA binding

Binding reactions as described above were assembled into individual wells of a 96-well black round bottom plate. The reactions were scanned using a plate reader with an excitation wavelength at 485nm and an emission wavelength of 528nm in the parallel and perpendicular direction (Pagano et al. 2011). The data were analyzed using Gen5 software to generate fluorescent polarization binding graphs. K_d_ measurements were calculated using Prism 9 software.

## ACKNOWLEDGEMENTS/Contributions

We thank Laura Vanderploeg for preparing the figures. We thank the National *Xenopus* Resource and Xenbase for valuable information and resources.

## Author contributions

Conceptualization: M.E.D., M.D.S.; Validation: M.E.D., T.P.T., M.D.S.; Formal analysis: M.E.D., M.D.S.; Investigation: M.E.D., T.P.T., M.D.S.; Writing - original draft: M.E.D., M.D.S.; Writing - review & editing: M.E.D., M.D.S., C.A.F; Supervision: M.D.S.; Project administration: M.D.S.; Funding acquisition: M.D.S.

## Supplemental Figure Legends

**Supplementary figure 1.**
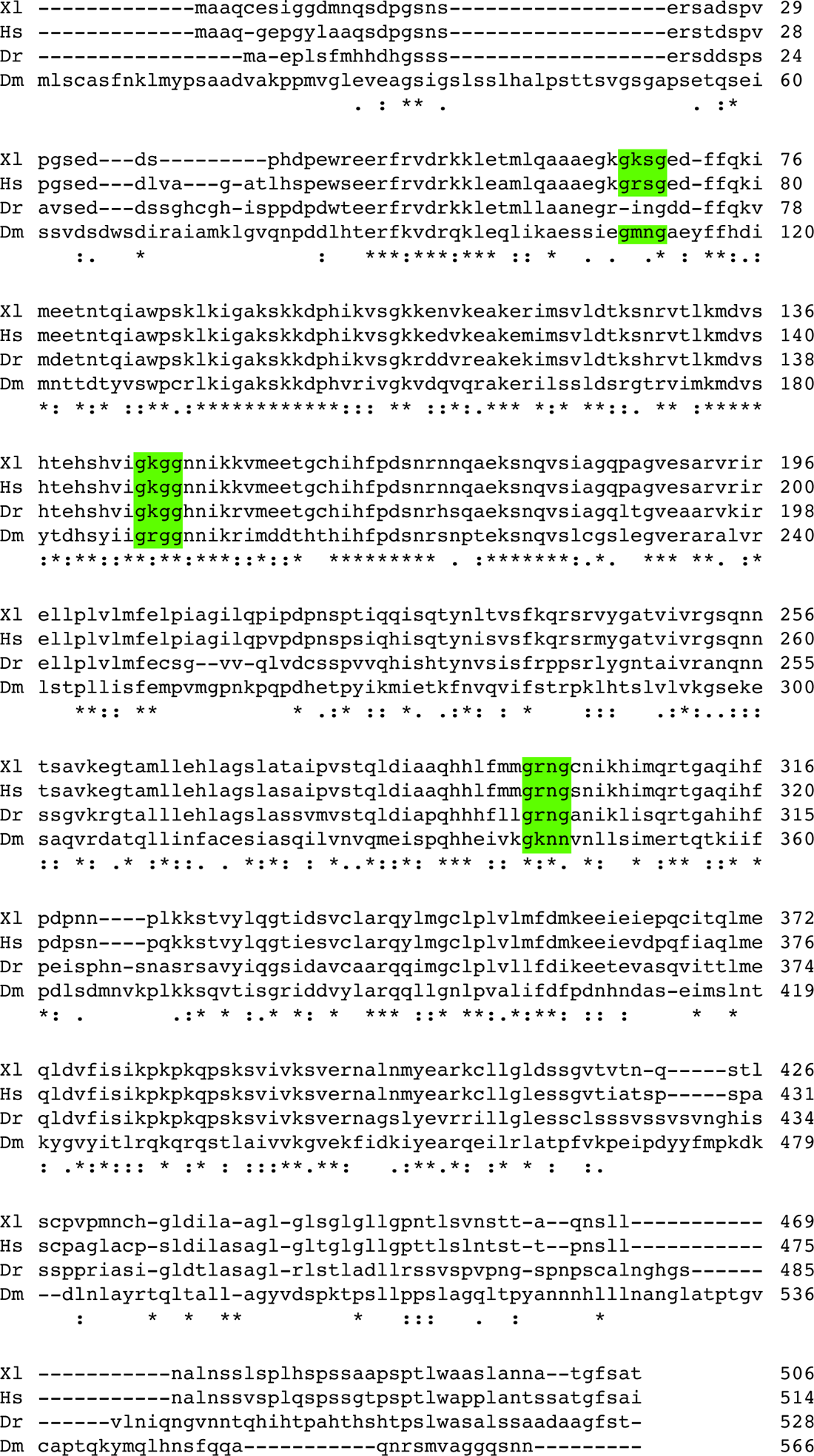
The evolutionarily conserved features of the N-terminal domain of Bicc1 proteins. Amino acid sequences from vertebrate and invertebrate Bicc1 proteins were analyzed with Clustal Omega. Each of the KH domains and the associated GXXG motifs are indicated.

**Supplementary figure 2.**
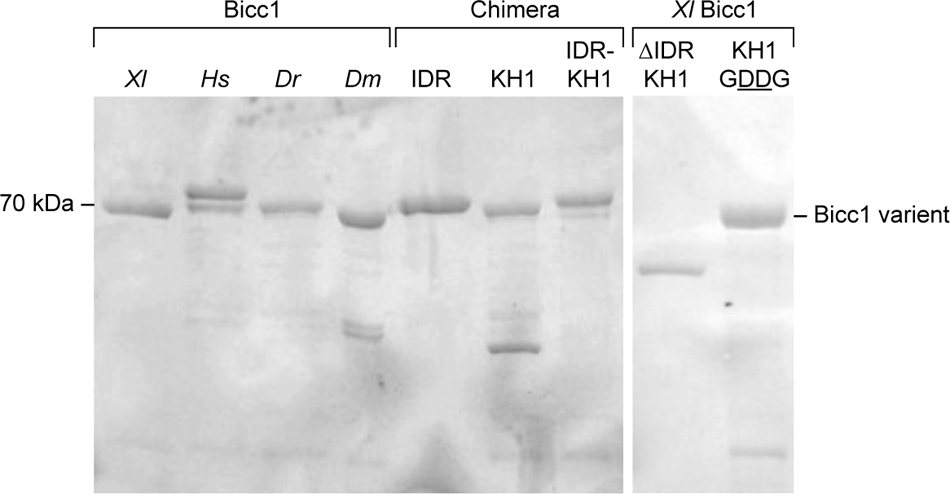
Bicc1 protein variants used for RNA binding assays. The different Bicc1 protein variants (aa 1-506) purified from *E.coli* and used for RNA binding assays were analyzed by SDS polyacrylamide gel electrophoresis.

**Supplementary figure 3.**
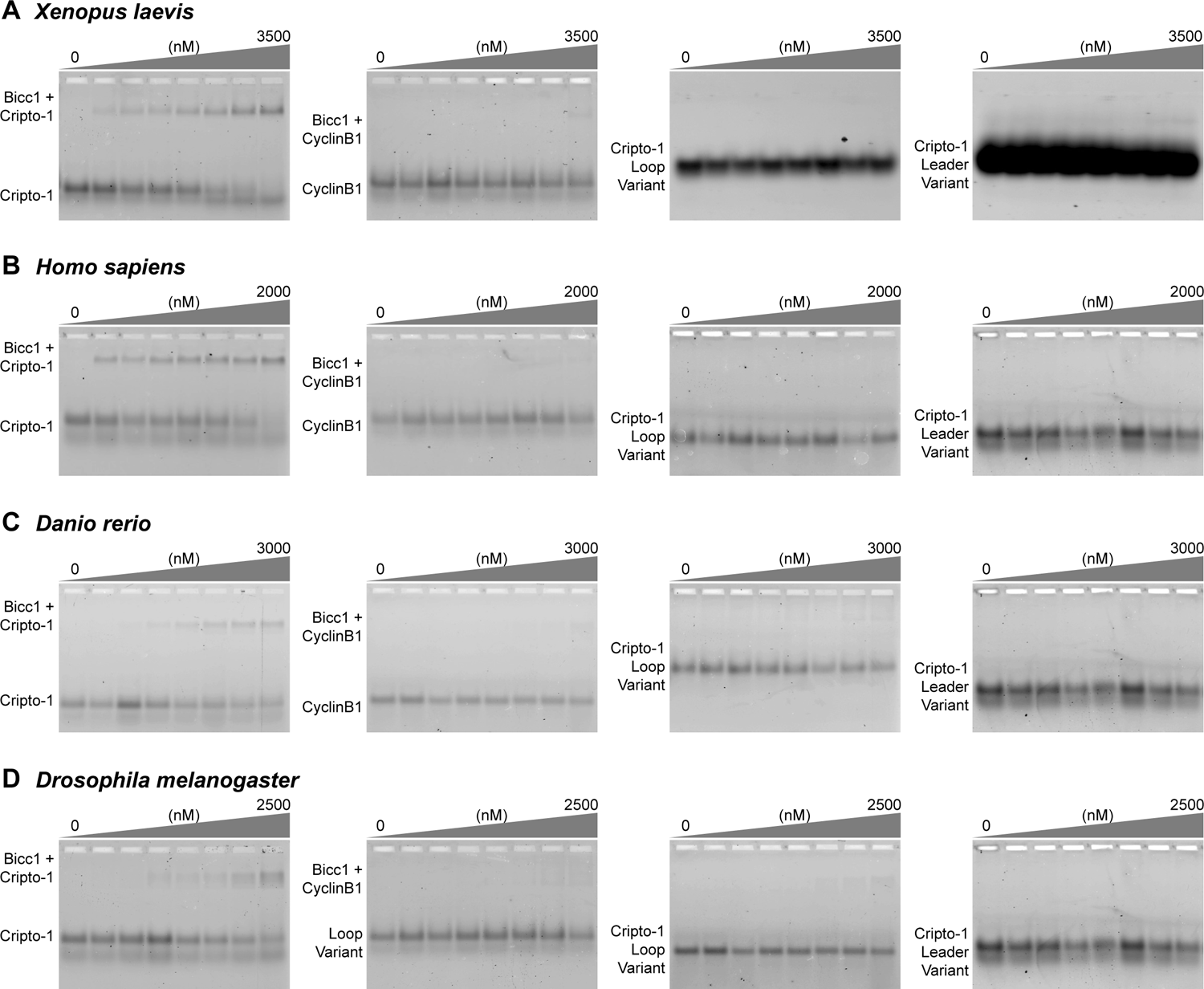
Electrophoretic mobility shift assay analysis of Bicc1 evolutionary variants. Titrations of increasing protein concentration of Bicc1 evolutionary variants *Xenopus laevis* **(A)**, *Homo sapiens* **(B)**, *Danio rerio* **(C)**, and *Drosophila melanogaster* **(D)** with RNA variants (from left to right): Wild-Type *Xl* Cripto-1 RNA, Cyclin B1 RNA, Cripto-1 Loop Variant, and Cripto-1 Leader variant.

**Supplementary figure 4.**
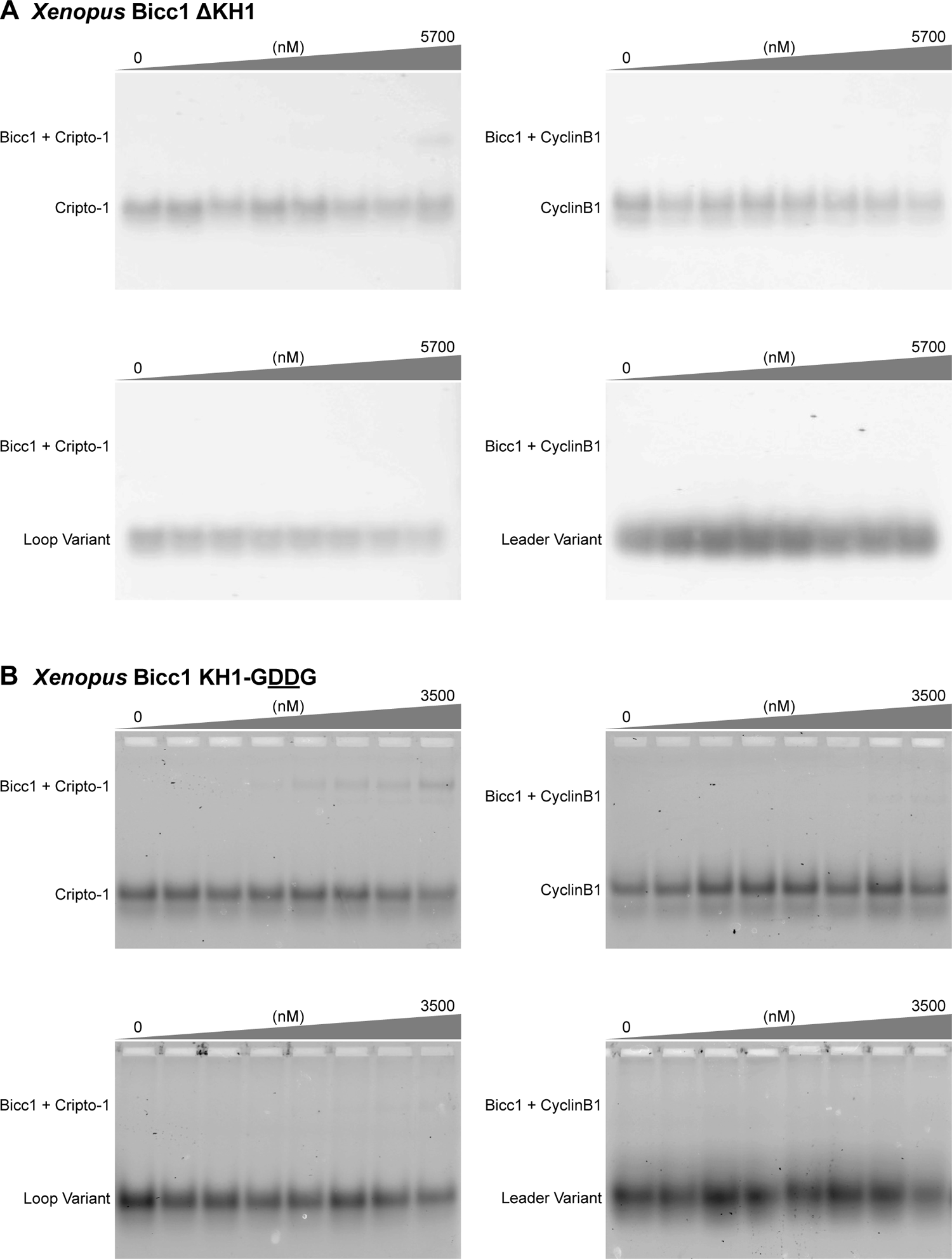
Electrophoretic mobility shift assay analysis of *Xl*Bicc1 KH1 variants. Titrations of increasing Bicc1 protein concentrations analyzed by EMSA of *Xenopus* Bicc1 ΔKH1 **(A)** and *Xenopus* Bicc1 KH1-GDDG mutant **(B)** with Wild-Type *Xl* Cripto-1 (top left), Cyclin B1 RNA (top right), Cripto-1 Loop Variant (bottom left), and Cripto-1 Leader variant (bottom right) RNA variants.

**Supplementary figure 5.**
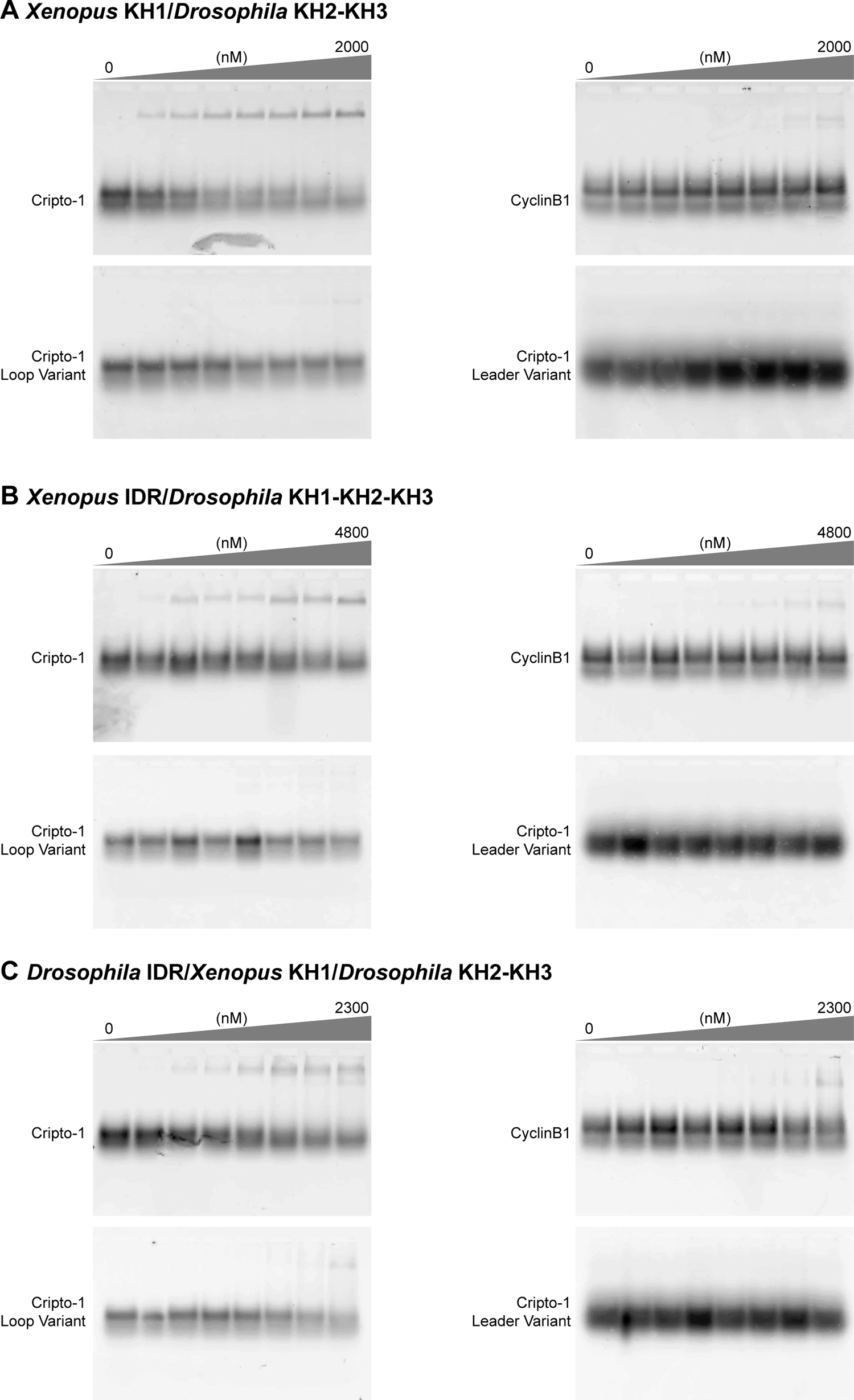
Electrophoretic mobility shift assay analysis of Bicc1 *Xenopus*-*Drosophila* fusion proteins. Titrations of increasing *Xenopus-Drosophila* protein concentrations analyzed by EMSA of *Xenopus* KH1*/Drosophila* KH2-KH3 **(A)**, *Xenopus IDR/Drosophila* KH1-KH2-KH3**, (B)** and *Drosophila IDR/Xenopus* KH1*/Drosophila* KH2-KH3 **(C)** with Wild-Type *Xl* Cripto-1 (top left), Cyclin B1 RNA (top right), Cripto-1 Loop Variant (bottom left), and Cripto-1 Leader variant (bottom right) RNA variants.

---

**Supplementary Table 1.**
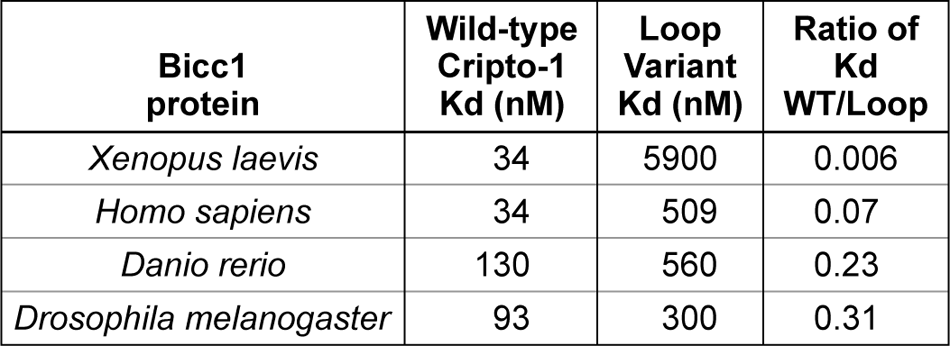
K_d_s determined from fluorescent polarization assays of the Bicc1 proteins for the wild-type and loop variant Cripto-1 RNAs as well as the ratio of wild-type K_d_ /Loop Variant K_d_.

**Supplementary Table 2.**
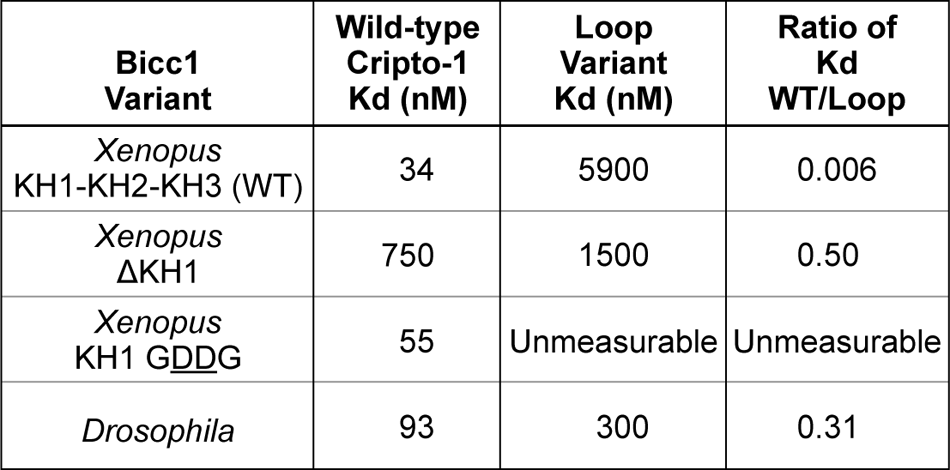
K_d_s of the *Xenopus* Bicc1 protein variants and *Drosophila* Bicc1 for the wild-type and loop variant Cripto-1 RNAs as well as the ratio of wild-type K_d,_ /Loop Variant K_d_.

**Supplementary Table 3.**
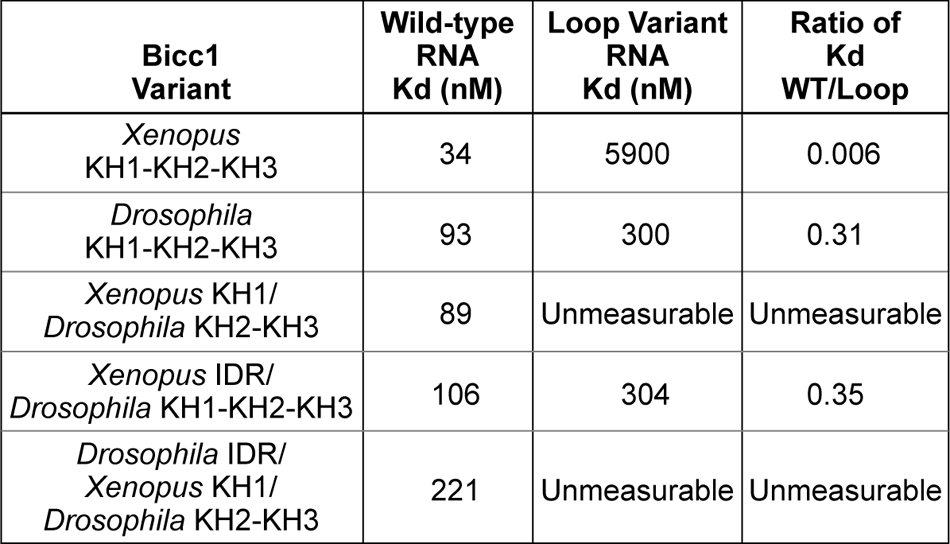
K_d_s of the *Xenopus* Bicc1 protein variants, *Drosophila* Bicc1 and the *Xl*KH1-*Dm*KH2KH3 chimeric protein for the wild-type and loop variant Cripto-1 RNAs as well as the ratio of wild-type K_d_ /Loop Variant K_d_.

## REFERENCES

1. Arai Y, Totoki Y, Hosoda F, Shirota T, Hama N, Nakamura H, Ojima H, Furuta K, Shimada K, Okusaka T, et al. 2014. Fibroblast growth factor receptor 2 tyrosine kinase fusions define a unique molecular subtype of cholangiocarcinoma. Hepatology 59: 1427–1434. doi:10.1002/hep.26890

2. Bermingham R, Carballedo A, Lisiecka D, Fagan A, Morris D, Fahey C, Donohoe G, Meaney J, Gill M, Frodl T. 2012. Effect of genetic variant in BICC1 on functional and structural brain changes in depression. Neuropsychopharmacology 37: 2855–2862. doi:10.1038/npp.2012.158

3. Biswas J, Patel VL, Bhaskar V, Chao JA, Singer RH, Eliscovich C. 2019. The structural basis for RNA selectivity by the IMP family of RNA-binding proteins. Nat Commun 10: 4440. doi:10.1038/s41467-019-12193-7

4. Bouvrette DJ, Price SJ, Bryda EC. 2008. K homology domains of the mouse polycystic kidney disease-related protein, Bicaudal-C (Bicc1), mediate RNA binding in vitro. Nephron Exp Nephrol 108: e27–34. doi:10.1159/000112913

5. Bouvrette DJ, Sittaramane V, Heidel JR, Chandrasekhar A, Bryda EC. 2010. Knockdown of bicaudal C in zebrafish (Danio rerio) causes cystic kidneys: a nonmammalian model of polycystic kidney disease. Comp Med 60: 96–106.

6. Bull AL. 1966. Bicaudal, a genetic factor which affects the polarity of the embryo in Drosophila melanogaster. Journal of Experimental Zoology 161: 221–241. doi:10.1002/jez.1401610207

7. Chao JA, Patskovsky Y, Patel V, Levy M, Almo SC, Singer RH. 2010. ZBP1 recognition of beta-actin zipcode induces RNA looping. Genes Dev 24: 148–158. doi:10.1101/gad.1862910

8. Cogswell C, Price SJ, Hou X, Guay-Woodford LM, Flaherty L, Bryda EC. 2003. Positional cloning of jcpk/bpk locus of the mouse. Mamm Genome 14: 242–249. doi:10.1007/s00335-002-2241-0

9. Cristinziano G, Porru M, Lamberti D, Buglioni S, Rollo F, Amoreo CA, Manni I, Giannarelli D, Cristofoletti C, Russo G, et al. 2021. FGFR2 fusion proteins drive oncogenic transformation of mouse liver organoids towards cholangiocarcinoma. J Hepatol 75: 351–362. doi:10.1016/j.jhep.2021.02.032

10. Davidson S, Shanley L, Cowie P, Lear M, McGuffin P, Quinn JP, Barrett P, MacKenzie A. 2016. Analysis of the effects of depression associated polymorphisms on the activity of the BICC1 promoter in amygdala neurones. Pharmacogenomics J 16: 366–374. doi:10.1038/tpj.2015.62

11. Dominguez D, Freese P, Alexis MS, Su A, Hochman M, Palden T, Bazile C, Lambert NJ, Van Nostrand EL, Pratt GA, et al. 2018. Sequence, Structure, and Context Preferences of Human RNA Binding Proteins. Mol Cell 70: 854–867.e9. doi:10.1016/j.molcel.2018.05.001

12. Dowdle ME, Imboden SB, Park S, Ryder SP, Sheets MD. 2017. Horizontal Gel Electrophoresis for Enhanced Detection of Protein-RNA Complexes. J Vis Exp 125: e56031. doi:10.3791/56031

13. Dowdle ME, Park S, Blaser Imboden S, Fox CA, Houston DW, Sheets MD. 2019. A single KH domain in Bicaudal-C links mRNA binding and translational repression functions to maternal development. Development 146: dev172486. doi:10.1242/dev.172486

14. Du Z, Lee JK, Fenn S, Tjhen R, Stroud RM, James TL. 2007. X-ray crystallographic and NMR studies of protein-protein and protein-nucleic acid interactions involving the KH domains from human poly(C)-binding protein-2. RNA 13: 1043–1051. doi:10.1261/rna.410107

15. Gamberi C, Lasko P. 2012. The Bic-C family of developmental translational regulators. Comp Funct Genomics 2012: 141386. doi:10.1155/2012/141386

16. Hollingworth D, Candel AM, Nicastro G, Martin SR, Briata P, Gherzi R, Ramos A. 2012. KH domains with impaired nucleic acid binding as a tool for functional analysis. Nucleic Acids Res 40: 6873–6886. doi:10.1093/nar/gks368

17. Lewis CM, Ng MY, Butler AW, Cohen-Woods S, Uher R, Pirlo K, Weale ME, Schosser A, Paredes UM, Rivera M, et al. 2010. Genome-wide association study of major recurrent depression in the U.K. population. Am J Psychiatry 167: 949–957. doi:10.1176/appi.ajp.2010.09091380

18. Lewis HA, Musunuru K, Jensen KB, Edo C, Chen H, Darnell RB, Burley SK. 2000. Sequence-specific RNA binding by a Nova KH domain: implications for paraneoplastic disease and the fragile X syndrome. Cell 100: 323–332. doi:10.1016/s0092-8674(00)80668-6

19. Maerker M, Getwan M, Dowdle ME, McSheene JC, Gonzalez V, Pelliccia JL, Hamilton DS, Yartseva V, Vejnar C, Tingler M, et al. 2021. Bicc1 and Dicer regulate left-right patterning through post-transcriptional control of the Nodal inhibitor Dand5. Nat Commun 12: 5482. doi:10.1038/s41467-021-25464-z

20. Mahone M, Saffman EE, Lasko PF. 1995. Localized Bicaudal-C RNA encodes a protein containing a KH domain, the RNA binding motif of FMR1. EMBO J 14: 2043–2055. doi:10.1002/j.1460-2075.1995.tb07196.x

21. Malakhov MP, Mattern MR, Malakhova OA, Drinker M, Weeks SD, Butt TR. 2004. SUMO fusions and SUMO-specific protease for efficient expression and purification of proteins. J Struct Funct Genomics 5: 75–86. doi:10.1023/B:JSFG.0000029237.70316.52

22. Minegishi K, Rothé B, Komatsu KR, Ono H, Ikawa Y, Nishimura H, Katoh TA, Kajikawa E, Sai X, Miyashita E, et al. 2021. Fluid flow-induced left-right asymmetric decay of Dand5 mRNA in the mouse embryo requires a Bicc1-Ccr4 RNA degradation complex. Nat Commun 12: 4071. doi: 10.1038/s41467-021-24295-2

23. Mohler J, Wieschaus EF. 1985. Bicaudal mutations of Drosophila melanogaster: alteration of blastoderm cell fate. Cold Spring Harb Symp Quant Biol 50: 105–111. doi:10.1101/sqb.1985.050.01.015

24. Mohler J, Wieschaus EF. 1986. Dominant maternal-effect mutations of Drosophila melanogaster causing the production of double-abdomen embryos. Genetics 112: 803–822. doi:10.1093/genetics/112.4.803

25. Nakel K, Hartung SA, Bonneau F, Eckmann CR, Conti E. 2010. Four KH domains of the C. elegans Bicaudal-C ortholog GLD-3 form a globular structural platform. RNA 16: 2058–2067. doi:10.1261/rna.2315010

26. Nicastro G, Candel AM, Uhl M, Oregioni A, Hollingworth D, Backofen R, Martin SR, Ramos A. 2017. Mechanism of β-actin mRNA Recognition by ZBP1. Cell Rep 18: 1187–1199. doi:10.1016/j.celrep.2016.12.091

27. Nicastro G, Taylor IA, Ramos A. 2015. KH-RNA interactions: back in the groove. Curr Opin Struct Biol 30: 63–70. doi:10.1016/j.sbi.2015.01.002

28. Ota KT, Andres W, Lewis DA, Stockmeier CA, Duman RS. 2015. BICC1 expression is elevated in depressed subjects and contributes to depressive behavior in rodents. Neuropsychopharmacology 40: 711–718. doi:10.1038/npp.2014.227

29. Pagano JM, Clingman CC, Ryder SP. 2011. Quantitative approaches to monitor protein-nucleic acid interactions using fluorescent probes. RNA 17: 14–20. doi:10.1261/rna.2428111

30. Park S, Blaser S, Marchal MA, Houston DW, Sheets MD. 2016. A gradient of maternal Bicaudal-C controls vertebrate embryogenesis via translational repression of mRNAs encoding cell fate regulators. Development 143: 864–871. doi:10.1242/dev.131359

31. Rothé B, Gagnieux C, Leal-Esteban LC, Constam DB. 2020. Role of the RNA-binding protein Bicaudal-C1 and interacting factors in cystic kidney diseases. Cell Signal 68: 109499. doi:10.1016/j.cellsig.2019.109499

32. Ryan J, Artero S, Carrière I, Maller JJ, Meslin C, Ritchie K, Ancelin M-L. 2016. GWAS-identified risk variants for major depressive disorder: Preliminary support for an association with late-life depressive symptoms and brain structural alterations. Eur Neuropsychopharmacol 26: 113–125. doi:10.1016/j.euroneuro.2015.08.022

33. Tran U, Zakin L, Schweickert A, Agrawal R, Döger R, Blum M, De Robertis EM, Wessely O. 2010. The RNA-binding protein bicaudal C regulates polycystin 2 in the kidney by antagonizing miR-17 activity. Development 137: 1107–1116. doi:10.1242/dev.046045

34. Wang Z, Zhou D, Li S, Zhang Y, Wang C. 2019. Underlying mechanisms of recombinant adeno-associated virus-mediated bicaudal C homolog 1 overexpression in the medial prefrontal cortex of mice with induced depressive-like behaviors. Brain Res Bull 150: 35–41. doi:10.1016/j.brainresbull.2019.05.008

35. Wessely O, De Robertis EM. 2000. The Xenopus homologue of Bicaudal-C is a localized maternal mRNA that can induce endoderm formation. Development 127: 2053–2062. doi:10.1242/dev.127.10.2053

36. Wessely O, Tran U, Zakin L, De Robertis EM. 2001. Identification and expression of the mammalian homologue of Bicaudal-C. Mech Dev 101: 267–270. doi:10.1016/s0925-4773(00)00568-2

37. Yang L, Wang C, Li F, Zhang J, Nayab A, Wu J, Shi Y, Gong Q. 2017. The human RNA-binding protein and E3 ligase MEX-3C binds the MEX-3-recognition element (MRE) motif with high affinity. J Biol Chem 292: 16221–16234. doi:10.1074/jbc.M117.797746

38. Ying X, Tu J, Wang W, Li X, Xu C, Ji J. 2019. FGFR2-BICC1: A Subtype Of FGFR2 Oncogenic Fusion Variant In Cholangiocarcinoma And The Response To Sorafenib. Onco Targets Ther 12: 9303– 9307. doi:10.2147/OTT.S218796

39. Zhang Y, Cooke A, Park S, Dewey CN, Wickens M, Sheets MD. 2013. Bicaudal-C spatially controls translation of vertebrate maternal mRNAs. RNA 19: 1575–1582. doi:10.1261/rna.041665.113

40. Zhang Y, Park S, Blaser S, Sheets MD. 2014. Determinants of RNA binding and translational repression by the Bicaudal-C regulatory protein. J Biol Chem 289: 7497–7504. doi:10.1074/jbc.M113.526426

